# Epidemiological and evolutionary consequences of CRISPR-Cas reactivity

**DOI:** 10.1101/2021.01.14.426534

**Authors:** Hélène Chabas, Viktor Müller, Sebastian Bonhoeffer, Roland R. Regoes

## Abstract

Adaptive immune systems face a control challenge: they should react with enough strength to clear an infection while avoiding to harm their organism. CRISPR-Cas systems are adaptive immune systems of prokaryotes that defend against fast evolving viruses. Here, we explore the CRISPR-Cas control challenge and look how its reactivity, i.e. its probability to acquire a new resistance, impacts the epidemiological outcome of a phage outbreak and the prokaryote’s fitness. We show that in the absence of phage evolution, phage extinction is driven by the probability to acquire at least one resistance. However, when phage evolution is fast, phage extinction is characterised by an epidemiological critical threshold: any reactivity below this critical threshold leads to phage survival whereas any reactivity above it leads to phage extinction. We also show that in the absence of autoimmunity, high levels of reactivity evolve. However, when CRISPR-Cas systems are prone to autoimmune reactions, intermediate levels of reactivity are evolutionarily optimal. These results help explaining why natural CRISPR-Cas systems do not show high levels of reactivity.

**Author summary:** CRISPR-Cas systems are adaptive immune systems that use a complex 3-step molecular mechanism to defend prokaryotes against phages. Viral infections of populations defending with CRISPR-Cas can result in rapid phage extinction or in medium-term phage maintenance. What controls phage fate? Using mathematical modeling, we show that two parameters control this outcome: the phage escape rate and CRISPR-Cas reactivity (i.e. its probability of resistance acquisition upon infection). Furthermore, CRISPR-Cas reactivity impacts host fitness. From this, we derive that 1) CRISPR-Cas reactivity is a key predictor of the efficiency and of the cost of a CRISPR-Cas system, 2) there is an optimal reactivity balancing the cost of autoimmunity and immune efficiency and 3) high phage escape rate selects for higher CRISPR-Cas reactivities.

## Introduction

The pressure viruses exert on their host has resulted in the evolution of various anti-viral immune defences, usually classified as innate or adaptive immunity. Adaptive immunity refers to systems that can acquire new specific targets of immunity during the first encounter with a parasite and memorise this resistance. To be efficient, these systems face a control challenge: they must lessen damage caused by the virus by limiting the spread of an infection and while avoiding dangerous autoimmune reactions (Bergstrom and Antia, 2006). This challenge has resulted in a tight regulation of their reactivity and any misregulation is harmful: lower reactivity results in a predisposition to severe infections whereas higher reactivity results in a propensity for autoimmunity.

The concept of adaptive immunity originated from the study of mammalian immune systems. However, it was later discovered that many others organisms carry some forms of adaptive immunity (Müller et al., 2018). In prokaryotes, adaptive immunity is represented by CRISPR-Cas systems (Clustered Regularly Interspaced Short Palindromic Repeats – CRISPR ASsociated) (Barrangou et al., 2007). These systems are composed of two loci: a CRISPR locus, that can be seen as a heritable library of small sequences derived from previously encountered viruses and a Cas locus that codes for all the proteins required for the system to work (Koonin et al., 2017). There are six known types of CRISPR-Cas systems that differ in their Cas genes composition, which can result in differences in their molecular mechanism (Koonin et al., 2017). However, the most common mechanism is as follows: when a cell is infected by a prokaryotic virus (phage), the invader can be detected by some Cas proteins and a sequence of approximatively 30-60 bp of its DNA (the protospacer) is integrated into the chromosomal CRISPR locus (where it is called a spacer): this is the acquisition step. Then, the CRISPR locus is transcribed and the spacer RNA is used as a guide to target the invader DNA and, upon matching, to trigger its degradation (expression and interference steps) (Barrangou et al., 2007; Hampton et al., 2020). As this mechanism relies on Watson-Crick pairing, viruses can escape by mutating their targeted protospacer (Deveau et al., 2008; Semenova et al., 2011; Sun et al., 2013).

Phages can evolve rapidly and, because for most CRISPR-Cas systems a single mutation is all that is needed to escape immunity, the escape from a spacer is usually fast (Deveau et al., 2008; Semenova et al., 2011; Sun et al., 2013). Recently, it was discovered that CRISPR-Cas systems prevent this through the generation of a diversity of spacers (van Houte et al., 2016; Chabas et al., 2018). This diversity of spacers arises during the acquisition step: phage genomes contain hundreds or thousands of potential protospacers and the choice among them is stochastic. As a consequence, each acquisition event leads to the acquisition of a different spacer in each prokaryotic cell, which at the population level, generates a diversity of spacers (Childs et al., 2012; Sun et al., 2013; van Houte et al., 2016; Paez-Espino et al., 2013; Heler et al., 2019). This diversity acts as an epidemiological shield against the spread of newly evolved escape mutants because when an escape mutant evolve against a given spacer, all hosts carrying another spacer still degrade this mutant, preventing its spread (Chabas et al., 2018). Overall, if diversity is high enough, this results in rapid phage extinction (van Houte et al., 2016).

The efficiency of CRISPR-Cas immunity (i.e. its ability to rapidly eradicate the phage population) therefore relies on its ability to generate diversity. The generation of diversity derives from the probability per infection that a cell acquires a random spacer (Heler et al., 2017), which we call CRISPR-Cas reactivity. However, CRISPR-Cas systems are also prone to autoimmunity i.e. to the acquisition of a spacer that targets a chromosomal sequence, which triggers its degradation and this is often thought to be lethal (Stern et al., 2010; Jiang et al., 2013; Vercoe et al., 2013; Gomaa et al., 2014; Wimmer and Beisel, 2020). Importantly, the level of autoimmunity of a CRISPR-Cas system is related to its reactivity (Levy et al., 2015; Workman et al., 2021). Therefore, it is likely that CRISPR-Cas systems face a similar control challenge as vertebrate adaptive immune systems: too much reactivity increases autoimmunity, but low levels of reactivity decrease the efficiency of the immune response (Weissman et al., 2020).

This control challenge is evidenced by three observations. First, CRISPR-Cas reactivity is low: one cell in a million for example for the *Streptococcus thermophilus* most active CRISPR-Cas system (Hynes et al., 2017). Second, scientists can easily generate CRISPR-Cas systems with higher reactivity but these are not the forms found naturally (Heler et al., 2017; Levy et al., 2015; Workman et al., 2021). Third, even though the study of CRISPR-Cas regulation is in its early stages, it is already clear that the reactivity of the system is tightly regulated (Patterson et al., 2017).

A previous theoretical work attempted to explore the CRISPR-Cas control challenge (Bradde et al., 2019). They found that autoimmunity decreases the optimal reactivity and that the optimal reactivity is governed by the interactions of the initial size of the bacteria and phages populations. However, in their work, they do not take into account that i) phages can infect bacteria carrying a spacer, which results in phage degradation (Barrangou et al., 2007), ii) spacers can be escaped by phage mutation (Deveau et al., 2008; Semenova et al., 2011; Sun et al., 2013), iii) CRISPR-Cas generates a diversity of resistance which protects the population against phage evolution (Childs et al., 2012; van Houte et al., 2016; Chabas et al., 2018). Consequently, it is unclear how a change in CRISPR-Cas reactivity changes the epidemiological outcome of a phage infection and the fitness of the prokaryote carrying it. Does an increase in CRISPR-Cas reactivity always improve its efficiency? Does a low reactivity always decrease bacterial fitness in the presence of phage? What is the impact of phage evolution on CRISPR-Cas efficiency and optimal reactivity? How does autoimmunity affect the optimal reactivity?

Answering these questions is experimentally challenging as it would require to finely modify CRISPR-Cas reactivity and phage mutation rates without altering other biological parameters. We therefore chose mathematical modeling to explore the consequences of CRISPR-Cas reactivity on the epidemiological outcome and on the bacterial fitness. Specifically, we developed a stochastic model of the coupled population dynamics and genetics that simulates the early dynamics of a virulent phage outbreak in a population carrying naive CRISPR-Cas immunity. Our model accounts i) for the ability of CRISPR-Cas to generate spacer diversity by the stochastic acquisition of spacers, ii) for phage escape of a given spacer through mutation and iii) for phage clearance resulting from the infection of cells carrying non-escaped spacers. In the absence of autoimmunity, we found that when phage evolution is slow, increasing CRISPR-Cas reactivity is beneficial as it increases the probability of generating at least one spacer that can target a phage. However, when phage evolution is fast, the survival of the phage population is governed by an epidemiological critical threshold: below a certain reactivity, the phage always remains in the system, whereas above this critical threshold, it is always driven to extinction. This is due to a non-linear relationship between CRISPR-Cas reactivity and the diversity of spacers that is generated. We also show that strains with higher reactivities have higher fitness. In the presence of autoimmunity, epidemiological outcomes are not modified, except for very high reactivities that impair bacterial survival. However, when strains with different reactivities compete, intermediate reactivities are selected in the presence of phages whereas the lowest reactivity is selected for in their absence. Finally, we show that lowering the propensity for autoimmunity (for example by the presence of a self/non-self discrimination mechanism) decreases its cost and therefore selects for higher levels of reactivity. We then discuss the implications of these findings for the evolutionary ecology of CRISPR-Cas and phages.

## Results

To study the relationship between CRISPR-Cas reactivity, spacer diversity and the outcome of phage infection, we develop a stochastic epidemiological model (see Methods for a full presentation of the model and Figure 1 for a graphic representation). Briefly, naive CRISPR-Cas bacteria are infected by virulent wildtype phage WT. Infection can lead to one of three outcomes with different probabilities: 1) the phage kills the bacterial cell and produces WT progeny, 2) the phage kills the cell and produces WT and a randomly-chosen mutated phage, 3) CRISPR-Cas kills the infecting phage by acquiring a randomly-selected spacer. WT phages can also enter a cell carrying a spacer and this results in phage death, without consequences for the cell. Escape phages can also infect a cell with a spacer and in this case, the outcome depends on the identity of the spacer and of the escape mutation. If the mutation escapes this specific spacer, the infection kills the cell and produces escape virions; if not, the phage is killed by the CRISPR-Cas system without consequences for the cell. These processes have been implemented stochastically and we can therefore follow the dynamics of each genotype, including its appearance and extinction (see Figure 2 for representative simulations).

**Figure 1:**
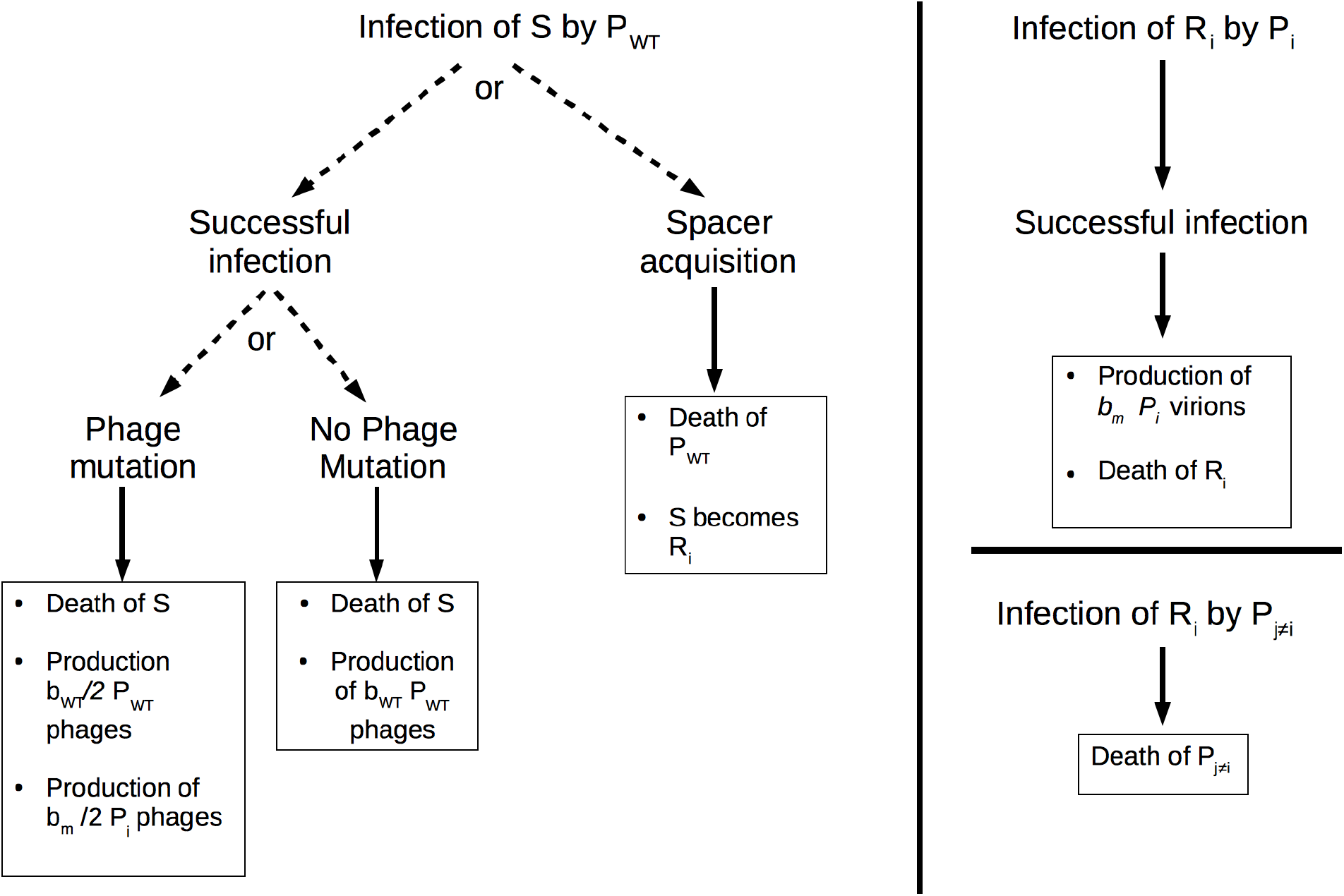
Important transitions in the model

**Figure 2:**
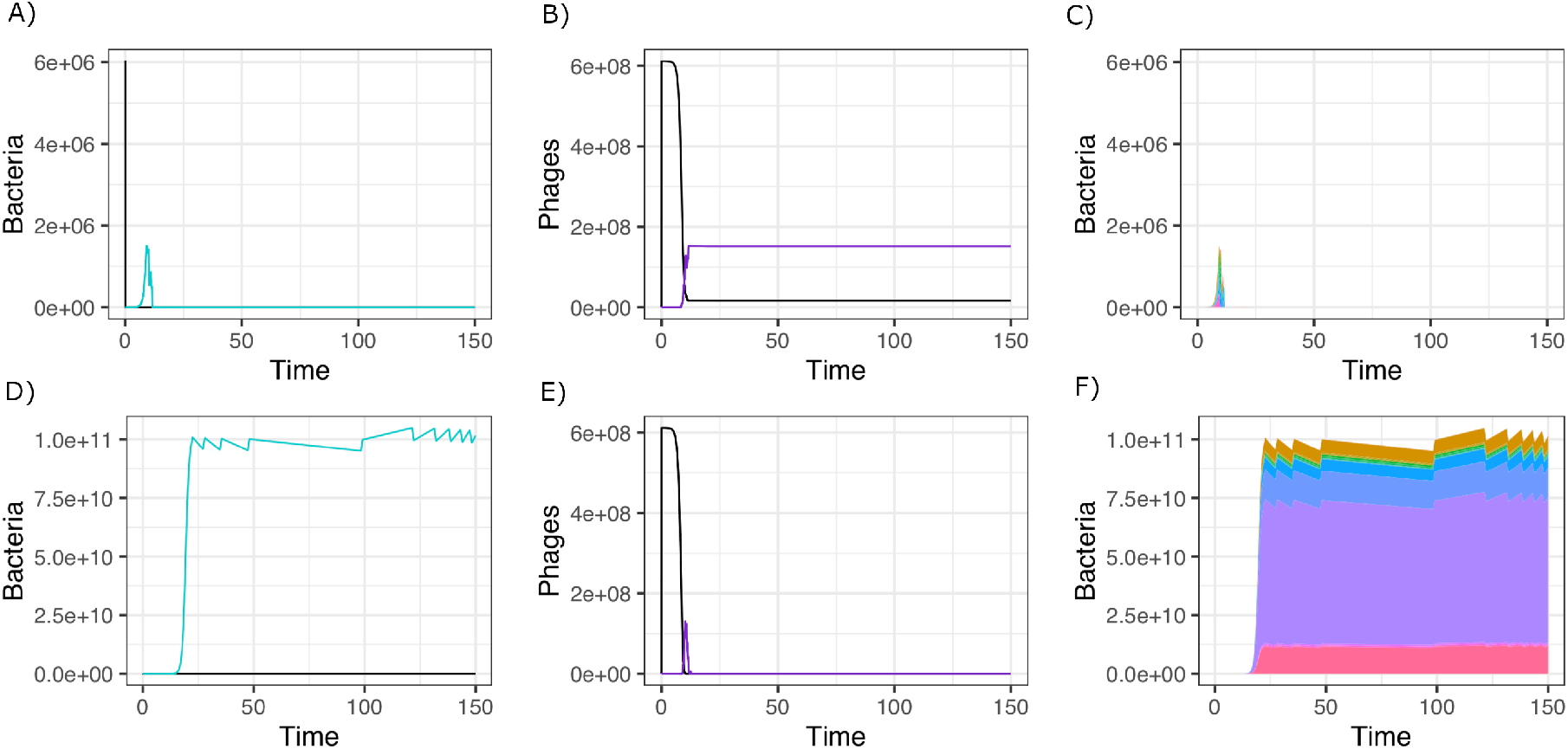
Example of time courses the model produces when 6 *\** 10^6^ S bacteria are infected by 10^5^ WT phages (*α* = 5.12 *\**10^*−*6^, *µ* = 10^*−*6^). In panel A, B and C, the phage drives the bacteria to extinction whereas in panel D, E and F the phage is driven to extinction by the CRISPR-Cas system. Panels A and D show the number of bacteria that are sensitive (black) and resistant (blue). Panels B and E show the number of phages that are WT (black) and escape mutants (purple). Panels C and F show the strains diversity of the bacterial population through time, each colour representing a strain.

### The diversity of spacers protects against phage epidemics

To evaluate the validity of our model, we check whether it predicts outcomes that have been experimentally observed. First, we test if the model predicts that initial spacer diversity protects the bacterial population against phage outbreaks. Indeed, it was shown experimentally that spacer diversity protects the bacterial population by preventing the spread of escape phages (van Houte et al., 2016). To do so, we run 100 simulations with an initial bacterial population composed of 50% naive CRISPR-Cas bacteria and 50% resistant cells with various levels of diversity. This population is infected by 10^5^ WT phages and we look at the probability of phage extinction at the end of the simulations. We find (Figure S1) that indeed, increasing the initial diversity of spacers in the host population increases the probability of phage extinction, which fits experimental results obtained by van Houte et al. (2016).

### Higher reactivities lead to higher diversity of spacers

Second, we want to confirm that higher reactivities lead to higher levels of spacers diversity. Indeed, Heler et al. found that a mutant with a higher CRISPR-Cas reactivity generates more diversity (Heler et al., 2017). We run 100 simulations of an outbreak due to an infection with 10^5^ WT phages that cannot evolve in a naive bacterial population. We look at the diversity of the bacterial population at the very beginning of the outbreak (at the time when sensitive bacteria are driven extinct). As expected, we observe that higher reactivities lead to higher levels of spacers diversity (Figure S2, panel A, black line). Notably, the relation between spacer diversity and CRISPR-Cas reactivity is non-linear. Instead, three phases can be observed: at low reactivity, the level of spacer diversity increases slowly; at intermediate reactivity, the diversity increases rapidly and at high reactivities, the system reaches the maximal number of potential spacers and saturates. Therefore, if any increases of spacer diversity are selected for, only increases in CRISPR-Cas reactivity that result in a transition from very low levels to intermediate levels of acquisition or an increase in the intermediate range can lead to an increase in bacterial spacers diversity.

### Phage survival depends on CRISPR-Cas reactivity and phage evolution

To understand the relationship between CRISPR-Cas reactivity and phage extinction, we run 100 simulations for various reactivities. For each, we calculate the probability of phage extinction by determining whether at least one of the phage genotype remains at the end of the simulation. We first look at the outcomes of outbreaks that are caused by phages that cannot escape CRISPR-Cas (*µ* = 0). We observe that when reactivity is low, rises in reactivity increase the probability of phage extinction (Figure 3, panel A, black line). When CRISPR-Cas reactivity reaches a certain value (around 10^*−*6^), the probability of phage extinction saturates to 1. Therefore, up to a certain value, evolving higher reactivities enhances the efficacy of CRISPR-Cas.

**Figure 3:**
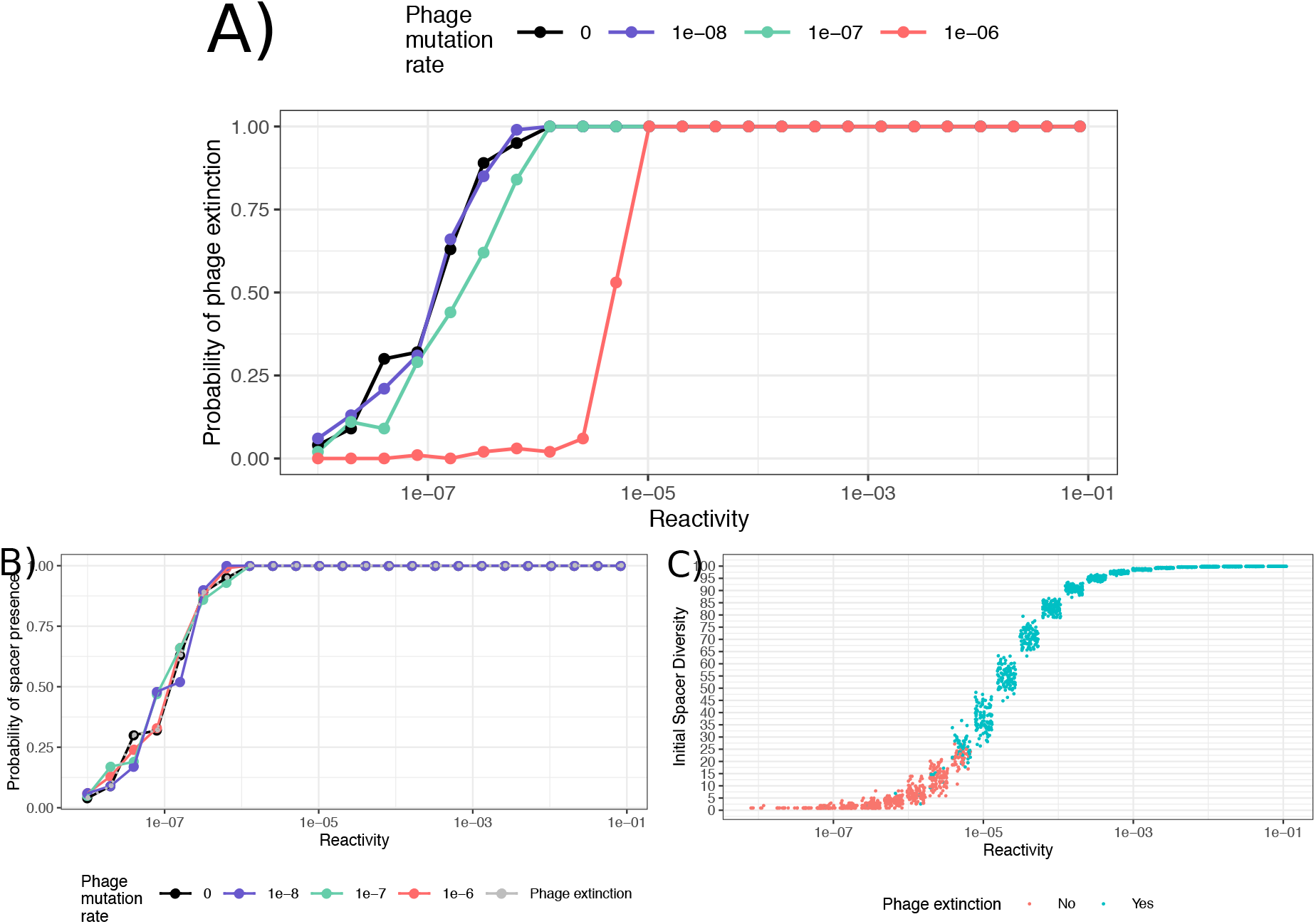
Influence of CRISPR-Cas reactivity on the probability of phage extinction in the absence of autoimmunity. A) Probability of phage survival when infecting bacteria with different reactivities. The different colours correspond to different levels of phage evolution (*µ*): black, no evolution (*µ* = 0), purple *µ* = 10^*−*8^; green *µ* = 10^*−*7^ and red *µ* = 10^*−*6^. B) Probability for bacteria with different reactivities to generate at least one single resistant cells. The different colours correspond to different levels of phage evolution (*µ*): in black, no evolution (*µ* = 0), in purple *µ* = 10^*−*8^; in green *µ* = 10^*−*7^ and in red *µ* = 10^*−*6^. The grey line corresponds to the probability of phage extinction in the absence of phage evolution *µ* = 0. C) Relationship between CRISPR reactivity (x-axis), initial spacer diversity (y-axis) and phage extinction (color) when phage evolution is high (*µ* = 10^*−*6^).

However in living systems, phages can escape CRISPR-Cas immunity by mutations. How does phage evolution change the outcome of an infection? We repeat the previous analysis for various probabilities of phage evolution. We find that when phages evolve slowly (*µ* = 10^*−*8^ and *µ* = 10^*−*7^), the situation does not massively differ from the case without evolution (Figure 3, panel A, purple and green lines). However, when phage evolution is fast (*µ* = 10^*−*6^), we observe that the outcome of the outbreak is determined by an epidemiological critical threshold: below a reactivity around 10^*−*5^, the probability of phage extinction is equal to zero, whereas above this critical threshold, phages are always driven to extinction by CRISPR-Cas immunity (Figure 3, panel A, red line). It is notable that phage evolution increases the value of the minimal CRISPR-Cas reactivity that ensures phage extinction: frequent phage evolution puts a high pressure on the CRISPR-Cas system.

As in our model, phage mutation results in a progeny containing both WT and escape virions (see Materials and Methods for details), we wonder if this assumption could alter this result. To test for this, we simulate outbreaks with phage mutation resulting in a progeny exclusively composed of one escape mutant and we find qualitatively the same results (Figure S3). In addition, in natural systems, the cost of a phage mutation can vary (Chabas et al., 2019) but it is fixed in our model. We wonder how a change in the cost of escape will alter the epidemiological outcome. To test for this, we repeat the same analysis removing or increasing (mutant burst size equals to 10% of the WT burst size) the phage escape cost. Qualitatively we find the same outcomes (Figure S4): only the slope and the value of the epidemiological threshold are modified.

In addition to purging viruses from the host population, immune systems also prevent the spread of a parasite. To see whether increased CRISPR-Cas reactivities decrease the spread of phages, we look at the size of phage outbreaks for 100 simulations. We find that increasing CRISPR-Cas reactivity decreases the size of the outbreak (Figure S5). Importantly, increasing CRISPR-Cas reactivity when it is too small to lead the phage to extinction, still results in a decrease in the size of the outbreak. As a consequence it seems beneficial for bacteria to evolve higher reactivities, even if these levels do not increase the probability of phage extinction.

### When phage evolution is slow, the probability of generating one single spacer drives the epidemiological outcome

How can we explain these results? First, we try to understand how in the absence of phage evolution, the increase in CRISPR-Cas reactivity results in a higher probability of phage extinction. We reason that if CRISPR-Cas reactivity is too low, there may be a non-zero probability that no bacteria acquire a spacer: then, the phage would spread in the population until all bacteria have been killed. To check for this, we looked at the number of simulations in which the proportion of resistant strains is positive at the beginning of the outbreak i.e. at the time when the sensitive cells go extinct. We find that indeed increasing CRISPR-Cas reactivity increases the probability of generating at least one resistant genotype (Figure 3, panel B). Critically, the relationship between CRISPR-Cas reactivity and the probability of phage extinction is identical to the one between CRISPR-Cas reactivity and the probability of generating at least one single resistant strain. This means that the probability of generating at least one resistant strain explains the impact of CRISPR-Cas reactivity on the probability of phage extinction.

### When phage evolution is fast, the probability to generate enough diversity explains the epidemiological outcome

When phage evolution is fast, the probability of generating one spacer does not recapitulate the epidemiological outcome. We wonder how can the epidemiological critical threshold be explained. We reason that spacer diversity is related to phage extinction and to CRISPR-Cas reactivity and that it is likely that phages are driven to extinction when initial spacer diversity is high. Because spacer acquisition and phage mutation are stochastic processes, each value of reactivity would then be associated with a probability to generate a certain diversity and each diversity in turn has a certain probability to drive the phages to extinction. This should result in simulations with low initial diversity having a low probability of phage extinction. Indeed, when we look at the initial spacer diversity depending on CRISPR-Cas reactivity and at the epidemiological outcome, we observe that simulations with low initial spacer diversity tend to result in phage survival, whereas simulations with high initial spacer diversity are likely to result in phage extinction (Figure 3, panel C). Importantly, there seems to be a diversity critical threshold: above this value, the phage is likely to go extinct whereas below this diversity phage extinction is unlikely.

### In the absence of autoimmunity, evolving higher reactivities is always beneficial

So far, we have looked at the impact of various CRISPR-Cas reactivities on the epidemiological outcome. What are the fitness consequences for the bacteria to evolve a higher CRISPR-Cas reactivity? To determine this, we simulate competition experiments. Two sensitive strains, that differ only in their reactivity, are mixed together and we infect this population with the phage *P*_*W T*_. At the end of the simulation, we calculate the proportion of each strain (sensitive + resistant cells) and we deduce the relative fitness. We observe that it is always beneficial for a strain to evolve higher reactivities (Figure 4, panel A). However, this is not what is observed in nature, where CRISPR-Cas reactivity is usually low (Hynes et al., 2017). In addition, scientists can easily generate CRISPR-Cas systems with higher reactivities (Heler et al., 2017; Levy et al., 2015; Workman et al., 2021), so the absence of strains with high reactivity is likely not the result of evolutionary constraints.

**Figure 4:**
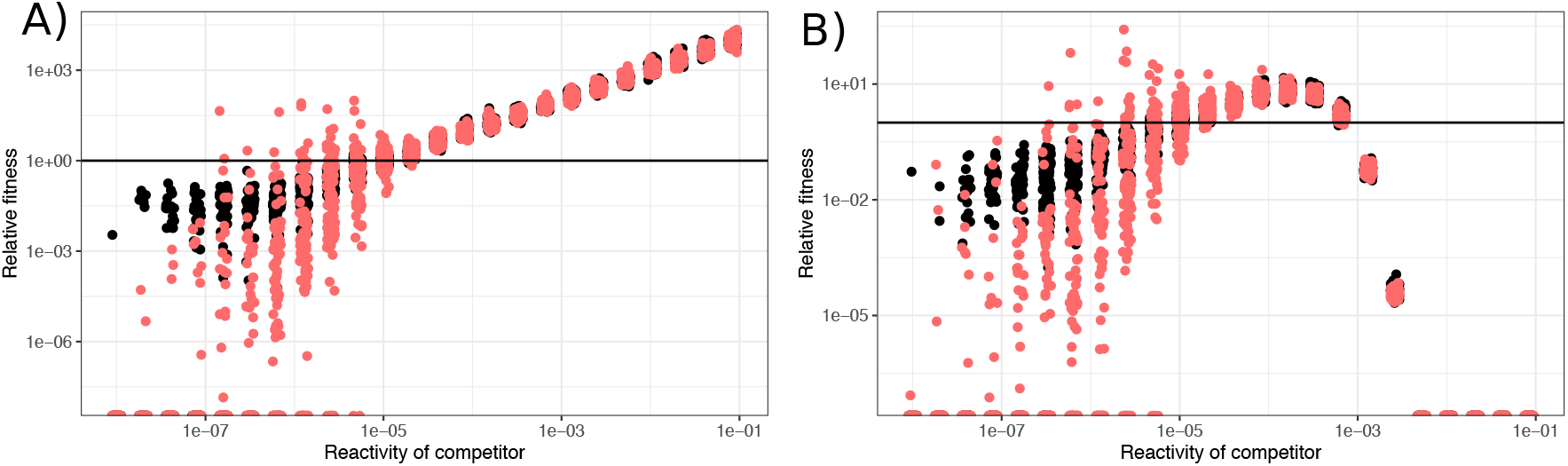
Fitness of a bacterial strain accross a range of reactivities when in competition with a strain with a reactivity *α* = 10^*−*5^. A) In the absence of autoimmunity, B) In the presence of autoimmunity (Propensity = 40). Colours correspond to phage evolution: in black, no evolution (*µ* = 0) and in red high phage evolution (*µ* = 10^*−*6^) Each point corresponds to one simulation. Only simulations in which bacteria survive the infection have been plotted.

### Autoimmunity selects for intermediate levels of reactivity

Natural CRISPR-Cas systems are prone to autoimmunity (Stern et al., 2010; Jiang et al., 2013; Vercoe et al., 2013; Gomaa et al., 2014), i.e. they can acquire spacers derived from the prokaryote chromosome. This autoimmunity has been shown to be linked to CRISPR-Cas reactivity and to decrease bacterial fitness (Heler et al., 2017; Workman et al., 2021). To explore the consequences of this autoimmunity, we add autoimmunity as an additional cause of bacterial death (see Methods for details).

Using this extended model, we look at the probability of phage extinction for various CRISPR-Cas reactivities. We find that the overall outcome of an outbreak is not modified by autoimmunity, except for very high reactivities, at which the bacteria are driven to extinction (Figure S6). Then we want to understand the evolutionary consequences of autoimmunity on bacteria: to do so, we simulate the competition between a WT strain with a reactivity of 10^*−*5^ and a mutant strain with various levels of reactivities. At the beginning of the simulation, both strains have equal represention. We observe that in the presence of autoimmunity, intermediate levels of reactivity are selected for (Figure 4, panel B). To get better insights into the cost of autoimmunity, we run competition in the absence of phages (Figure S7). We observe that it is always beneficial to have a lower reactivity, i.e. that autoimmunity always causes a detectable cost. In addition, the cost on fitness increases when autoimmunity increases: as a consequence, very high reactivity are massively selected against, whereas low and intermediate levels of reactivity are slightly deleterious.

### CRISPR-Cas propensity for autoimmunity determines the optimal reactivity

We reason that various CRISPR-Cas systems may not have the same propensity for autoimmunity and consequently the cost of autoimmunity can vary independently of the reactivity. To explore the consequences for bacterial fitness of various propensities, we run 100 competition simulations for various levels of propensity. We observe that the higher the propensity, the lower the optimal reactivity (Figure 5). Interestingly, we observe that high levels of propensity decreases the optimal reactivity at the level of the minimal reactivity needed to assures phage extinction. We wonder then if high levels of propensity can impair the efficiency of CRISPR-Cas immunity. To test for this, we ran 100 simulations of the infection of a bacterial population with high propensity to autoimmunity (*Propensity* = 4000) and look at phage survival. We find that indeed high levels of propensity makes really high levels of reactivity unbearable, however, the effect is not strong enough to prevent phage extinction for intermediate levels of reactivity (Figure S8).

**Figure 5:**
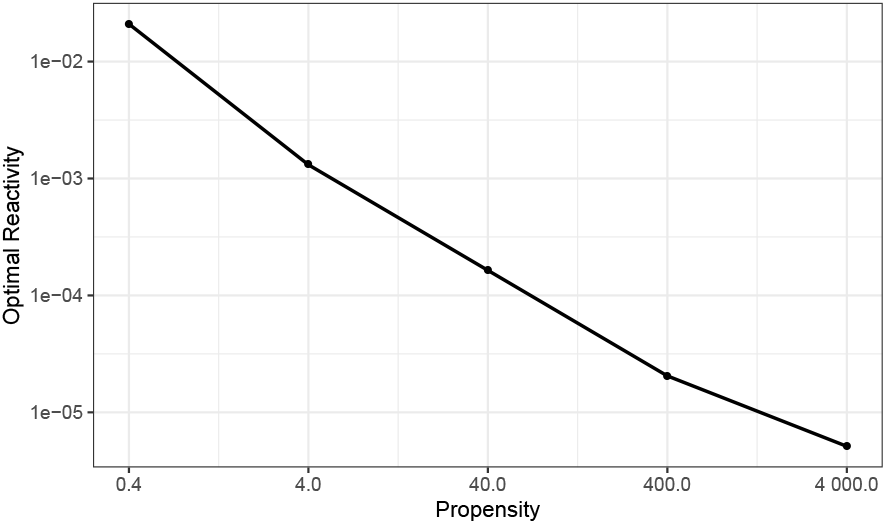
High levels of propensity selects for lower optimal reactivity. A control strain with a reactivity at 10^*−*5^ competes with a strain of interest, with various reactivity levels, in the presence of phage that can escape spacers (*µ* = 10^*−*6^). For each propensity (0.4, 4, 40, 400 and 4000), we took the median fitness of the bacteria and reported as the optimal reactivity the reactivity associated with the highest fitness.

### The evolution of surface mutants does not modify CRISPR-Cas optimal reactivity

Prokaryotes can defend against virulent phages using a wide variety of resistance strategies (Labrie et al., 2010; Goldfarb et al., 2015; Ofir et al., 2018; Doron et al., 2018; Millman et al., 2020). Especially, the evolution of surface mutants, i.e. cells that have modified, hidden or removed their phage receptor, is a frequent strategy of defence against phages (Gurney et al., 2019). We wonder how this alternative defence strategy impacts the optimal reactivity of CRISPR-Cas. To answer this question, we extend our model and let the sensitive population evolve surface mutations. The population of surface mutants cannot be infected by any phage genotype but it carries a fitness cost in the form of a reduced growth rate. We start by running competition simulations between two sensitive populations, both having the ability to defend either by surface modification or by CRISPR-Cas. These two strains only differ in their CRISPR-Cas reactivity: the control strain has a reactivity of 10^*−*5^ whereas the reactivity of strain of interest ranges from 10^*−*8^ to *≈* 8.10^*−*2^. The rate of surface mutants evolution can be low (*µ*_*SM*_ = 10^*−*6^) or high (*µ*_*SM*_ = 10^*−*4^). At the end of each simulation, we calculate the relative fitness of i) the strain of interest, ii) its CRISPR-Cas resistant genotypes and iii) surface mutants that have originated in the two sensitive populations. We find that CRISPR-Cas immunity explains most of the fitness of the prokaryote strain (with the exception of some simulations at very low CRISPR-Cas reactivity and high rates of surface mutants evolution) (Figure S9). We hypothesize that CRISPR-Cas optimal reactivity would not be modified by the possibility for surface mutations. To test for this, we determine, as previously described, the optimal reactivity of CRISPR-Cas systems with various propensities for autoimmunity. We find that the possibility to evolve surface mutants does not change the optimal reactivity of the CRISPR-Cas system (Figure S9).

## Discussion

We study the control challenge faced by the prokaryotic adaptive immune system, CRISPR-Cas. We find that the outcome of a phage outbreak is governed by CRISPR-Cas reactivity: 1) when resistance escape is impossible, rises in CRISPR-Cas reactivity increase the probability of phage extinction and this is governed by the probability for CRISPR-Cas to generate at least one single resistant genotype; 2) when the infected phage has a high rate of resistance escape, phage extinction is controlled by an epidemiological critical threshold: any value below a certain level of reactivity leads to phage persistence whereas any reactivity above leads to phage extinction. Importantly, this critical threshold is above the minimal reactivity that assures phage extinction in the absence of phage evolution. We also find that in the absence of autoimmunity, evolving higher reactivities is always beneficial. However, CRISPR-Cas susceptibility for autoimmunity results in the selection for intermediate reactivities: indeed, during an outbreak, high reactivity levels results in high fitness costs that negate the benefits of CRISPR-Cas. Finally, we show that the optimal reactivity level depends on the propensity of the system for autoimmunity.

In this work, we simulate outbreaks of two types of phages: phages that have a high rate of escaping spacers and phages that cannot escape spacers. This can correspond to two types of CRISPR-Cas immunity. Indeed, even if all types of CRISPR-Cas rely on Watson-Crick pairing to guide the CRISPR complex, their sensitivity to phage mutations varies greatly. For type I and type II systems, a single mutation in the protospacer adjacent motif or in the seed sequence is all that is needed to completely escape the spacer and this usually happens rapidly (Deveau et al., 2008; Chabas et al., 2018, 2019): therefore the model with fast phage evolution can be used to study these CRISPR-Cas systems. On the other hand, the interference mechanism of type III CRISPR-Cas systems is resilient to phage mutation and phage escape is extremely rare (Manica et al., 2013; Pyenson et al., 2017): the model in the absence of phage evolution can therefore be used to study interactions between phages and type III CRISPR-Cas systems. Our model predicts important differences in the epidemiological outcomes for type I/II and type III CRISPR-Cas systems. For type I/II CRISPR-Cas systems, the control of phage evolution is mediated by spacer diversity and there is a reactivity critical threshold whereas for type III CRISPR-Cas systems, spacer diversity does not impact phage extinction and the epidemiological outcome is driven by the probability to generate at least one resistant genotype. Because the minimal reactivity that ensure to acquire at least one spacer is lower than the reactivity critical threshold, this would suggest that type III CRISPR-Cas systems evolve lower reactivities than type I/II CRISPR-Cas systems. This lower reactivity for type III would indeed give an advantage to the cell as it would minimise the cost of autoimmunity.

Experimental data to support these predictions are lacking. Even if the importance of spacer diversity for the efficiency of type I/II CRISPR-Cas systems has been documented both experimentally and theoretically (van Houte et al., 2016; Chabas et al., 2018; Payne et al., 2018), the relationship between phage extinction probability and CRISPR-Cas reactivity has, as far as we know, not been explored. In addition, to the best of our knowledge, there have been no experimental studies of the coevolution of phages and naive type III CRISPR-Cas system. This is most likely due to the lack of type III CRISPR-Cas systems showing naive spacer acquisition under laboratory conditions and this makes the discovery of naive acquisition by the type III CRISPR system of *Thermus thermophilus* very promising (Artamonova et al., 2020).

In this work, we show that during a virulent phage outbreak, CRISPR-Cas autoimmunity selects for intermediate levels of reactivity (Figure 4). We also show that in the absence of phages, evolving higher reactivity is always costly (Figure S7), which selects for CRISPR-Cas systems with the lowest reactivity. As the presence or the absence of phages changes the selection pressure on the optimal reactivity, we expect that the optimal reactivity of CRISPR-Cas systems to be lower in environments where phage outbreaks are rare than in environments where they are frequent. One way for CRISPR-Cas systems to respond to this change in selection is to have their reactivity tightly regulated. Studies of the regulation of CRISPR-Cas show that CRISPR-Cas expression is finely regulated (Patterson et al., 2017). For example, it was shown that in some bacteria, CRISPR-Cas reactivity is upregulated by quorum sensing and this has been explained by the higher risk of phage outbreaks when cell density is high (Høyland-Kroghsbo et al., 2013; Patterson et al., 2016; Høyland-Kroghsbo et al., 2017). This finding makes sense in the light of our predictions as in the absence of phage infection, autoimmunity makes the system always costly and therefore CRISPR-Cas reactivity should evolve towards a minimum when the risk of infection is low.

From a phage perspective, we show that lower reactivities increase phage survival. Consequently, it is probably beneficial for phages to decrease CRISPR-Cas reactivity. Many phages carry anti-CRISPR proteins, i.e. small proteins that inhibit some CRISPR-Cas systems (Borges et al., 2017). To the best of our knowledge, most of them inhibit interference (Wiegand et al., 2020), a handful inhibit both acquisition and interference (Vorontsova et al., 2015) and none inhibit only acquisition: our model suggests that families of anti-CRISPR that decrease CRISPR-Cas reactivity would be beneficial.

Finally, one should not forget that an observed reactivity is also influenced by the ecological conditions: for example, it has been proposed that infections at lower temperature boosts CRISPR-Cas reactivity by slowing down the intra-host phage kinetics and therefore letting more time for the system to react (Høyland-Kroghsbo et al., 2018). This calls for caution when measuring and studying of CRISPR-Cas reactivity as the value measured might be the result of the system itself, its regulation, the infective phage and ecological conditions and may have limited prediction power if any of this changes.

It is striking that our model predicts an epidemiological critical threshold that is around 10^*−*5^, ten times higher than the reactivity of *S. thermophilus*’ most active CRISPR-Cas system (Hynes et al., 2017). Importantly, when infected by a virulent phage, *S. thermophilus*, defending exclusively with its CRISPR-Cas systems, survives and lead the phage to extinction in the long term (Paez-Espino et al., 2013; Common et al., 2019). There are two potential hypotheses that may explain this discrepancy between theoretical predictions and experimental data. First, the epidemiological outcome may depend on the value of the parameters. While we carefully parameterized our model using all available estimates from the experimental literature, some key parameters have, to our knowledge, not been estimated yet. Most notably, there is no estimate of phage infectivity. To test if our particular choice for the infectivity is the reason for the discrepancy, we ran the model with various infectivities. Varying the value of the infectivity only marginally affected the results (see supplementary figure S10). Therefore, we are confident that this is unlikely to explain this discrepency. Second, we have been making simplifying assumptions. Importantly, contrary to our model, CRISPR systems can acquire multiple spacers against a single phage and this is likely important for phage extinction. In addition, many CRISPR-Cas systems display some form of priming, which boosts the acquisition of a second spacer, either because the cell already carries a partially matching spacer (type I) or because it already possesses an efficient spacer (type II) (Datsenko et al., 2012; Fineran et al., 2014; Nussenzweig et al., 2019). The impact of these on the relationship between CRISPR-Cas reactivity, the epidemiological outcome, autoimmunity and prokaryote fitness remain to be explored and it is possible that this would change the epidemiological outcome for a given reactivity and the value of the optimal reactivity.

How can we experimentally assess the validity of our model assumptions and predictions? Testing the model predictions and assumptions requires a model system for which the CRISPR-Cas reactivity can be finely tuned. In our view, the most promising system for such work is the *Streptococcus pyogenes* type II CRISPR-Cas system, as its reactivity can be modified by mutations of the Cas9 protein and/or of the long-form tracrRNA (Heler et al., 2017; Workman et al., 2021). Therefore, mutants with various reactivities can be challenged by a virulent phage. As our model only looks at the short term dynamics, such infections should last less than 5 days. At the end of the experiments, phage extinction/survival can be detected by a stamping assay. Given the stochasticity of the outbreak, such an experiment would have to be reproduced sufficiently often to allow the precise quantification of the probability of phage extinction. Such a protocol would assess the existence of the epidemiological critical threshold. In addition, it would also be possible to study the relationship between CRISPR-Cas reactivity, genetic diversity and phage extinction/survival by sequencing the CRISPR array of randomly chosen replicates. Using sequencing data, the spacer diversity can be calculated and the relationship between spacer diversity, reactivity and phage extinction can be assessed. Finally, by knocking-out the nuclease function of Cas9, it is possible for such system to block interference while simultaneously conserving spacer acquisition (Wei et al., 2015; Jinek et al., 2012). Therefore, one can experimentally study the relationship between CRISPR-Cas reactivity and its level of autoimmunity. To do so, mutants with various reactivities can have their interference function knocked-out and be grown in the absence of phages. After deep-sequencing of their CRISPR-Cas array, the rate of autoimmunity of each strain can be calculated and variations in autoimmunity levels can be compared to variations in reactivities. As we propose to use a type II CRISPR-Cas system, rapid phage escape is expected. We think that a similar approach using a type III CRISPR-Cas system can be used to test the model predictions when phage escape is impossible.

## Material and Methods

### Model definition

In our mathematical model, we consider a bacterial population composed of a sensitive strain *S* carrying a naive CRISPR-Cas system, i. e. with no pre-exisiting spacers. Upon infection by phage *P*_*WT*_, *S* can acquire a spacer with a probability *α* and evolve resistance against *P*_*WT*_. We assume that the phage genome possesses *n*_*s*_ protospacers and that the CRISPR-Cas system randomly acquires one of them with equal probability. As a result, the infection of *S* will lead to the evolution of a diversified population composed of a subset of resistant strains *R*_*i*_, all carrying a single different spacer (=resistance) against phage WT. All these strains follow a logistic growth with a growth rate *g* and a total carrying capacity *K*.

However, the infection of *S* by *P*_*WT*_ can also be successful because the CRISPR-Cas system fails to stop the infection and the phage reproduces on *S*. A successful infection can either lead to the amplification of *P*_*WT*_ and produce *b*_*WT*_ progeny phages, or result in the escape of the phage from one spacer. Specifically, each protospacer has a probability *µ* to mutate and escape the spacer that targets it. The impact of a mutation on the fraction of the progeny carrying the mutation depends on the phage mode of replication (linear, binary, intermediate). Some phages are known to have a mode of replication that is close to binary whereas others have one close to linear (Sanjuán et al., 2010) and to the best of our knowledge, the replication mode of the majority of phages is unknown. Therefore, here, we assume an intermediate state where each phage genotype produces half of their respective progeny: 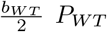 and 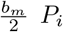. Each of the escape phage *P*_*i*_ can successfully infect the corresponding resistant bacteria *R*_*i*_ in addition to the sensitive strain *S*. If they infect another resistant strain *R*_*j*_, they are degraded by the CRISPR-Cas system with no consequences for the bacteria. Note that *b*_*m*_ *< b*_*WT*_: this results in escape phage mutants *P*_*i*_ having a lower fitness than *P*_*WT*_, an observation that has been made recently (Chabas et al., 2019).

The interaction between bacterial and phage populations described above can be summarized by the following differential equations:

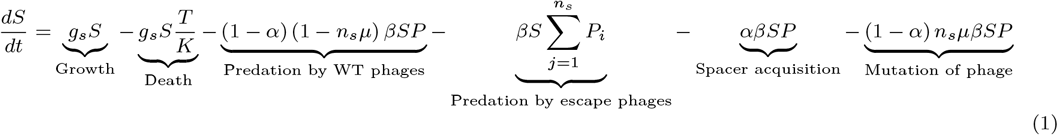

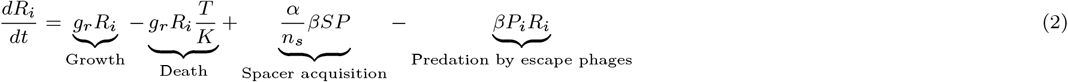

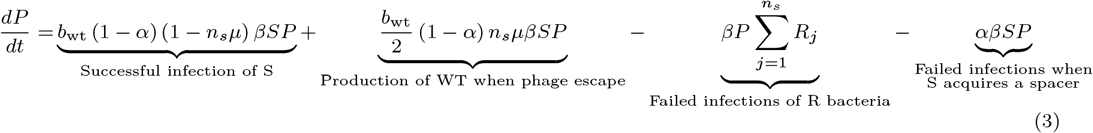

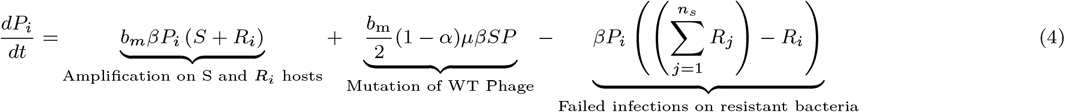

We implemented the dynamics described by Equations 1–4 stochastically. A list of all model variables and parameters and their definition can be found in table 1. Figure 1 illustrates the different processes we considered in our model.

**Table 1:** Summary of the variables and parameters used in the model.

| Variables | Biological interpretation |  |
| --- | --- | --- |
| $S$ | Population size of sensitive bacteria (no spacer) | |
| $R_i$ | Population size of resistant bacteria carrying spacer i | |
| $T$ | Sum of all bacterial population sizes | |
| $P$ | Population size of WT phage (no escape mutation) | |
| $P_i$ | Population size of escape phage (escape mutation in protospacer i) | |
| Parameters | Biological interpretation | Value |
| $\alpha$ | CRISPR-Cas reactivity: probability to acquire 1 spacer | variable |
| $\mu$ | Mutation probability of a protospacer | variable |
| $g$ | Bacterial growth rate per hour | $1.3/h$ |
| $K$ | Bacterial carrying capacity | $10^{11}$ |
| $n_s$ | Number of phage protospacers | 100 |
| $\beta$ | Infectivity rate constant | $10^{-6}/h$ |
| $b_{WT}$ | Burst size of the WT phage ($P$) | 100 |
| $b_m$ | Burst size of the escape phages ($P_i$) | 70 |
| *Propensity* | CRISPR-Cas propensity to autoimmunity | variable |
| $\mu_{SM}$ | Mutation rate leading to surface mutants | $10^{-6}$ |
| $g_{SM}$ | Growth rate of surface mutants | $1/h$ |

We are aware that our model does not take into consideration some complexities of CRISPR-Cas biology, such as priming (Fineran et al., 2014), acquisition of multiple spacers (Barrangou et al., 2007), heterogeneity in the probability of choosing a spacer (Heler et al., 2019; Modell et al., 2017), or multiple phage mutations. In addition, we also did not add a natural phage decay: indeed, each cell can only acquire one spacer, an event that occurs within 24 hours (Barrangou et al., 2007; Deveau et al., 2008; Cady et al., 2012). Consequently, we study outcomes occuring in several days, a timeframe where natural phage decay can be neglected in laboratory conditions.

Overall, our model is similar to the model developed by Childs et al. (2012).

### Mathematical modelling of autoimmunity

To test the impact of autoimmunity on bacterial fitness and CRISPR-Cas immunity, we added autoimmunity as an additional cause of bacterial death. To the best of our knowledge, the precise relationship between CRISPR-Cas reactivity and CRISPR-Cas autoimmunity is unknown. However, we know that an increase in reactivity results in higher autoimmunity (Levy et al., 2015; Workman et al., 2021). In addition, there is limited evidence that an increase in reactivity is proportional to autoimmunity (Levy et al., 2015). We therefore modelled the rate of death due to autoimmunity as being proportional to CRISPR-Cas reactivity and to the number of bacteria. In addition, the cost of autoimmunity can be modulated by a parameter representing the propensity of CRISPR-Cas for autoimmunity, independently of its reactivity: biologically, this can be understood as the propensity for the bacterial chromosome to be targeted by CRISPR-Cas, for example because of the density of the bacterial chromosome in potential protospacers or in sequences inhibiting acquisition (like Chi sites (Levy et al., 2015)) or because the bacterial CRISPR-Cas system possesses an efficient self/non-self distinction mechanism.

The rate of death due to autoimmunity is therefore implemented as the following term that is subtracted from Equation 1 and 2:

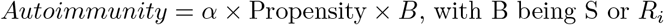

### Mathematical modelling of surface mutants evolution

To test for the impact of an alternative resistance strategy on CRISPR-Cas optimal reactivity, we extended our model by adding an additional population, SM, which is generated from the S population at a rate *μ*_*SM*_ * *S*. These cells follow a logistic growth with a rate *g*_*SM*_ *< g* and they are not infected by any of the phage genotypes.

### Bacterial spacer diversity

As a quantitative measure of the bacterial spacer diversity, we calculated the Simpson index from the frequency of each bacterial genotype *p*_*i*_:

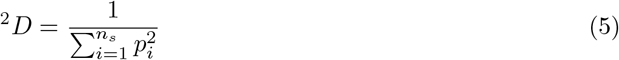

### Fitness calculation

We calculate relative fitness of two competing bacterial strains from their initial and final frequencies, *p*_*i*_ and *p*_*f*_ using the following formula:

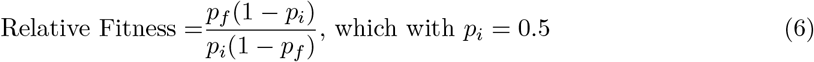

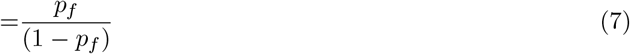

If the two strains are going extinct, relative fitness is set to 0. If the control strain goes extinct (i.e. *p*_*f*_ = 1), the relative fitness is equal to “Inf".

### Model implementation

All simulations and analyses were implemented and conducted in the language for statistical computing R 3.6.3 (R Core Team, 2018) and the library tidyverse (1.3.0) (Wickham et al., 2019). To implement the model (Equation 1–4) stochastically, we used the Gillespie algorithm in the package <u>adaptivetau</u> (Johnson, 2019).

## Acknowledgements

The authors thank Judith Bouman and Peter Ashcroft for their help in coding, Thomas Aubier and Mircea Sofonea for scientific discussions and Bruce Levin, Brandon Berryhill and Rodrigo Garcia Gonzales for valuable feedbakcs on an earlier version of this manuscript. This work was supported by an ETH Zurich Postdoctoral Fellowship (https://ethz.ch/en/research/research-promotion/eth-fellowships.html) to HC. VM was supported by the ELTE Thematic Excellence Programme 2020 funded by the National Research, Development and Innovation Office of Hungary (TKP2020-IKA-05). The funders had no role in study design, data collection and analysis, decision to publish, or preparation of the manuscript.

## Authors contributions

Conceptualization: HC; Methodology: HC, VM, RRR; Investigation: HC; Writing - Original Draft: HC; Writing - Review & Editing: HC, VM, SB, RRR; Visualization: HC; Supervision: SB, RRR; Project Administration: SB; Funding Acquisition: HC.

## Code Availability

All codes will be made available on github at the time of publication and have been submitted as supplementary informations for review purposes.

