## Supplemental figures for "Epidemiological and evolutionary consequences of CRISPR-Cas reactivity"

### Supplementary informations – Epidemiological and Evolutionary Consequences of CRISPR-Cas Reactivity

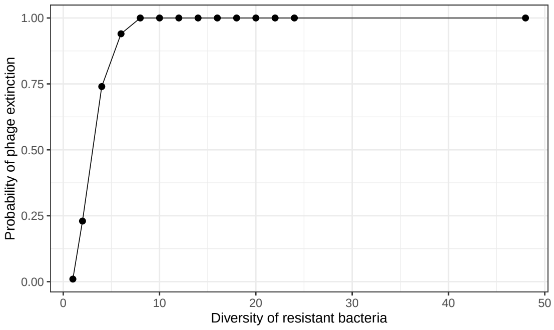


Figure S1: Impact of initial diversity on the probability of phage extinction.
The figure shows the probability of phage extinction in 100 simulations. At the beginning of the simulation, the bacterial population was composed of $6 * 10^6$ bacteria with an equal representation of sensitive bacteria (S) and different numbers of resistant bacterial genotypes.

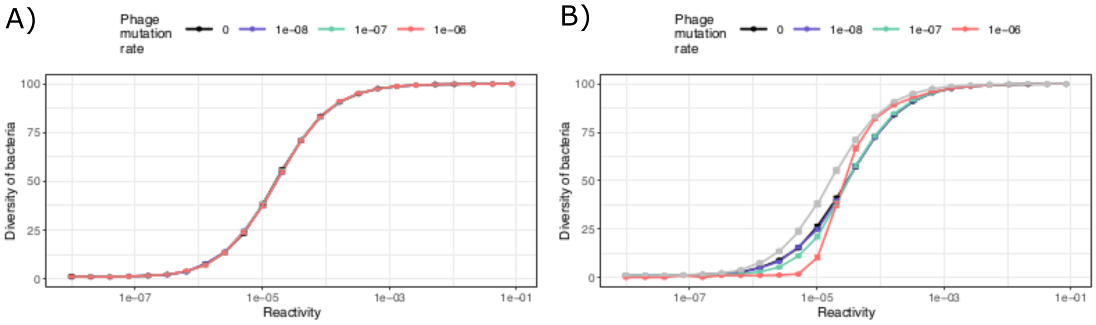


Figure S2: Influence of CRISPR reactivity on the mean diversity of newly generated spacers at the beginning (when S goes extinct, panel A) or at the end (panel B) of the outbreak.
Simulations that resulted in bacterial extinction are excluded from the calculation. The black curve represents the initial diversity of spacers where phage cannot evolve ($\mu = 0$) and the purle, green and red curves when phages can evolve ($\mu = 10^{-8}$, $\mu = 10^{-7}$, $\mu = 10^{-6}$ respectively). On Panel B, the grey line represents the initial diversity. Error bars correspond to 95% confidence intervals.

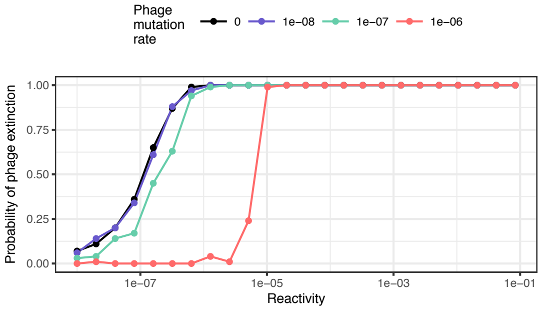


Figure S3: Probability of survival, for a phage infecting bacteria with various reactivities when phage mutation results in a progeny exclusively composed of escape mutants.

The different colours corresponds to different levels of phage evolution ($\mu$): black, no evolution ($\mu = 0$), purple $\mu = 10^{-8}$; green $\mu = 10^{-7}$ and red $\mu = 10^{-6}$.

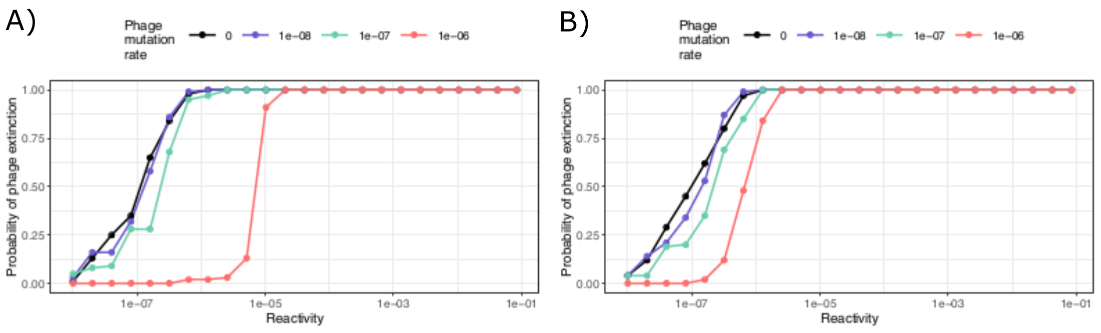


Figure S4: Influence of the cost of escaping CRISPR-Cas on the probability of phage extinction in the absence of autoimmunity.

Probability of phage survival when infecting bacteria with various reactivities. The different colours corresponds to different levels of phage evolution ($\mu$): black, no evolution ($\mu = 0$), purple $\mu = 10^{-8}$; green $\mu = 10^{-7}$ and red $\mu = 10^{-6}$. A) No cost, B) High fitness cost (burst size of mutants equals to 10% of phage WT burst size.)

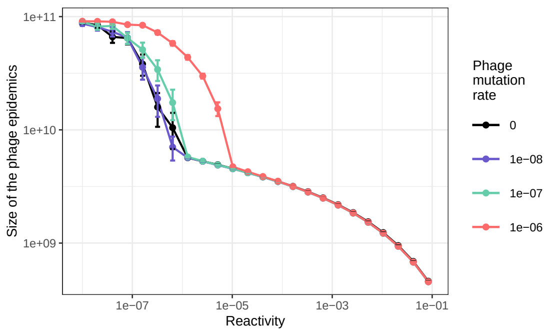


Figure S5: Influence of CRISPR reactivity on the size of a phage outbreak infecting bacteria using CRISPR immunity.

The colors represent phage evolution: black, no evolution ($\mu = 0$); purple, green and red $\mu = 10^{-8}$, $\mu = 10^{-7}$, $\mu = 10^{-6}$ respectively. Error bars corresponds to 95% confidence intervals.

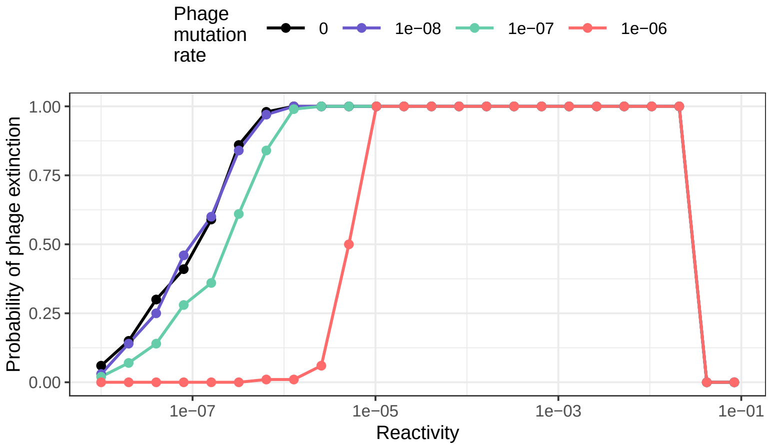


Figure S6: Influence of CRISPR reactivity on the probability of phage extinction in the presence of autoimmunity.

The different colors corresponds to different levels of phage evolution ($\mu$): in black, no evolution ($\mu = 0$), in purple $\mu = 10^{-8}$; in green $\mu = 10^{-7}$ and in red $\mu = 10^{-6}$.

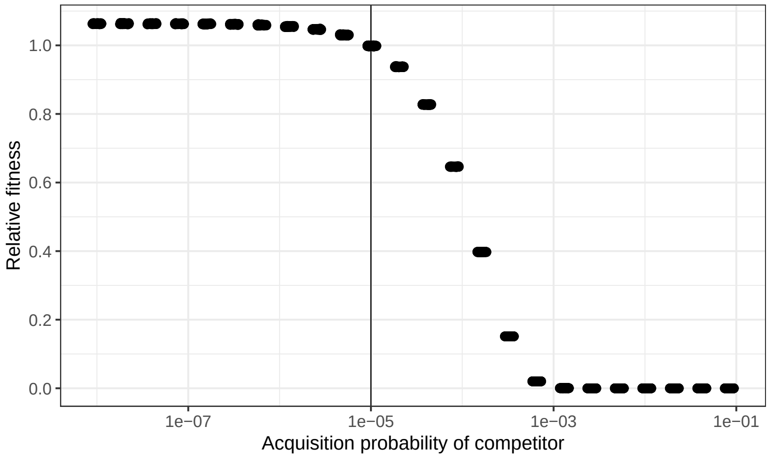


Figure S7: Fitness of a bacteria with various reactivity competing against a strain with a reactivity $\alpha = 10^{-5}$.

Each competition has been simulated 100 times and for each of them, the relative fitness have been plotted.

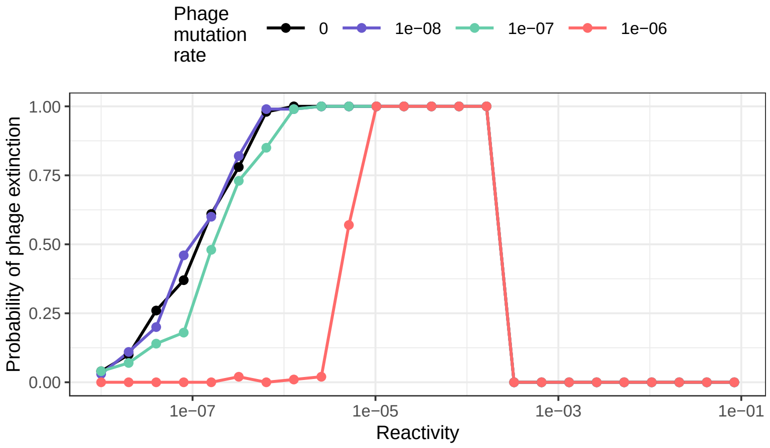


Figure S8: Influence of CRISPR reactivity on the probability of phage extinction with high propensity for autoimmunity (4000).

Probability of phage survival when infecting bacteria with various reactivities. The different colors corresponds to different levels of phage evolution ($\mu$): in black, no evolution ($\mu = 0$), in purple $\mu = 10^{-8}$; in green $\mu = 10^{-7}$ and in red $\mu = 10^{-6}$.

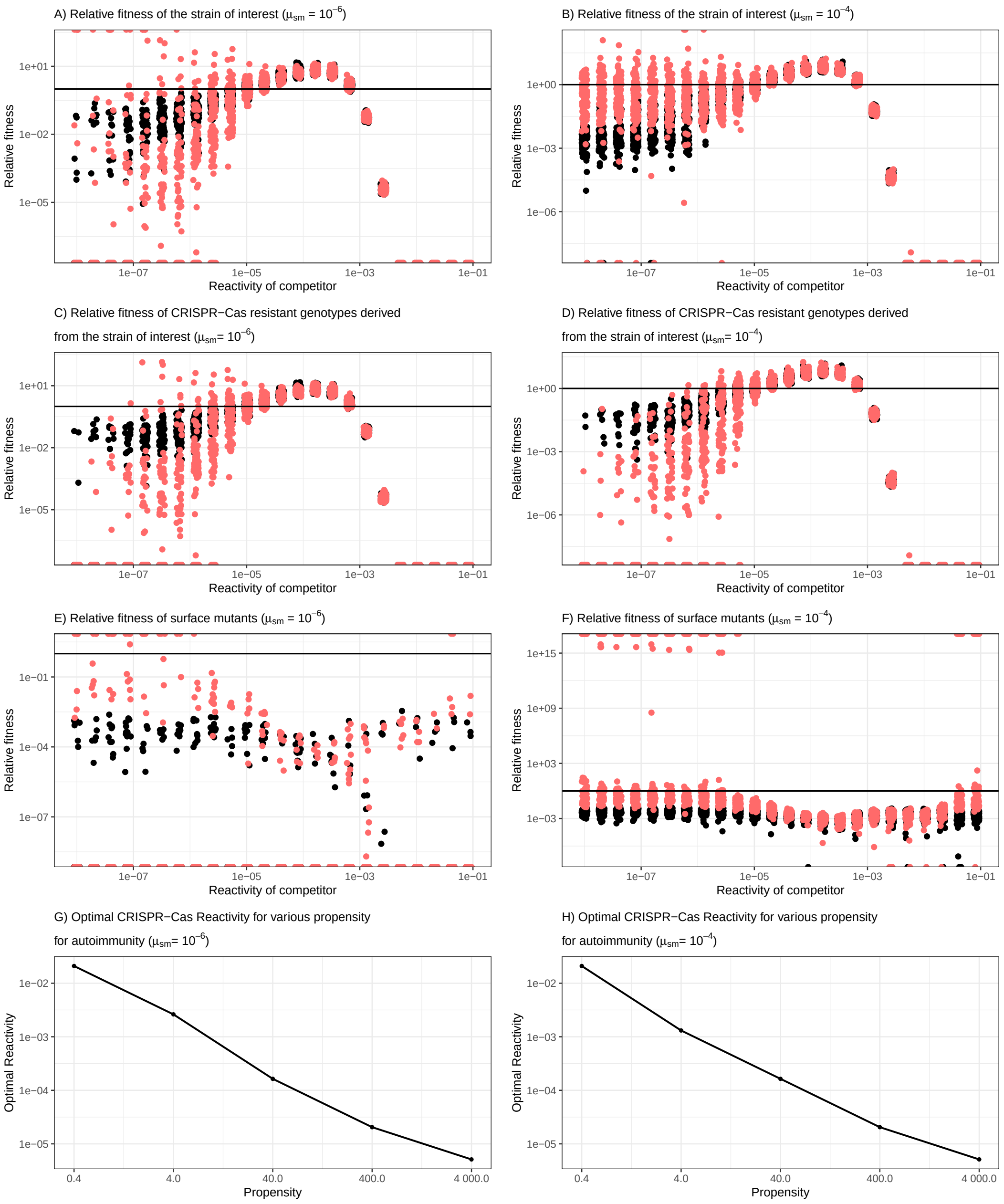


Figure S9: The evolution of surface mutants does not change CRISPR-Cas optimal reactivity.

Relative fitness of A,B) the strain of interest; C,D) the CRISPR-Cas resistant genotypes derived from it and E,F) surface mutants, at the end of a competition between a control strain (with a reactivity of $10^{-5}$) and a strain of interest (various reactivity levels), in the presence of phage that cannot (black) or can (red, $\mu = 10^{-6}$) escape CRIPSR-Cas immunity. The evolution of surface mutants is either rare ($\mu_{SM} = 10^{-6}$) (A,C and E) or frequent ($\mu_{SM} = 10^{-4}$) (B,D and F). G,H) Optimal reactivity of a strain of interest with various reactivity competing against a reference strain with a reactivity of $10^{-5}$ in the presence of a phage that can escape spacers ($\mu = 10^{-6}$) when surface mutant evolution is rare (G) or frequent (H). Both strains can also evolve surface modifications. For each propensity (0.4, 4, 40, 400 and 4000), we took the median fitness of the bacteria and reported as the optimal reactivity the reactivity associated with the highest fitness.

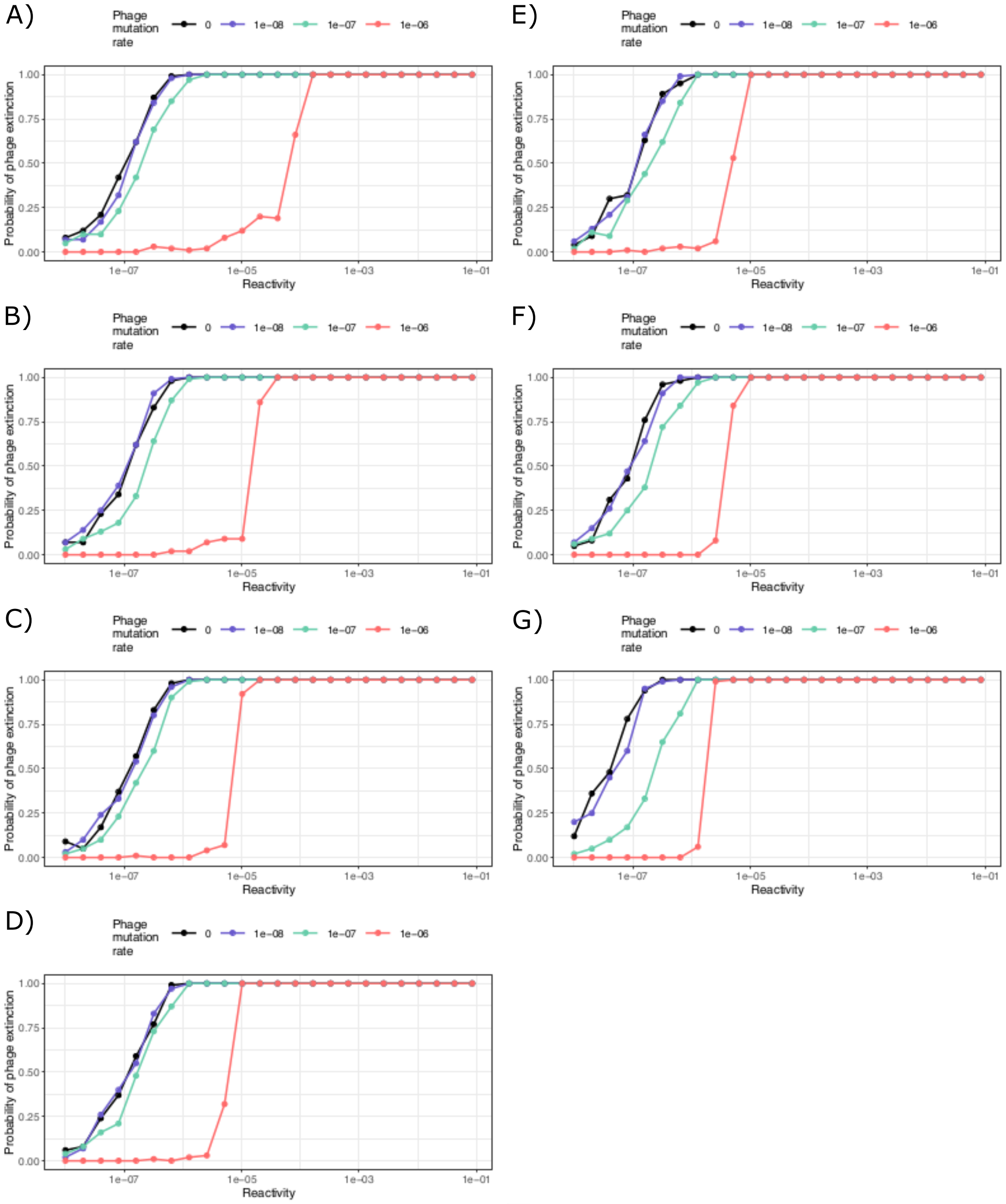


Figure S10: Influence of phage infectivity on the epidemiological outcome in the absence of autoimmunity.

Probability of phage survival when infecting bacteria with various reactivities. The different colors corresponds to different levels of phage evolution ($\mu$): in black, no evolution ($\mu = 0$), in purple $\mu = 10^{-8}$; in green $\mu = 10^{-7}$ and in red $\mu = 10^{-6}$. The different panels represent various phage infectivity: A) $\beta = 10^{-2}$, B) $\beta = 10^{-3}$, C) $\beta = 10^{-4}$, D) $\beta = 10^{-5}$, E) $\beta = 10^{-6}$, F) $\beta = 10^{-7}$, G) $\beta = 10^{-8}$
